# The evolution of a placenta is not linked to increased brain size in poeciliid fishes

**DOI:** 10.1101/2020.11.07.372615

**Authors:** P. K. Rowiński, J. Näslund, W. Sowersby, S. Eckerström-Liedholm, B. Rogell

**Affiliations:** Department of Zoology, Stockholm University, 106 91 Stockholm, Sweden; Department of Aquatic Resources, Institute of Freshwater Research, Swedish University of Agricultural Sciences, Stångholmsvägen 2, 17893 Drottningholm, Sweden; Department of Biology, Faculty of Science, Osaka City University, 3-3-138 Sugimoto Sumiyoshi-ku, Osaka 558-8585, Japan

**Keywords:** maternal provisioning, maternal effects, placenta, matrotrophy, expensive tissue, brain allometry, trade-offs, poeciliids

## Abstract

Maternal investment is considered to have a direct influence on the size of energetically costly organs, including the brain. In placental organisms, offspring are supplied with nutrients during pre-natal development, potentially modulating brain size. However, the coevolution of the placenta and brain size remains largely unknown in non-mammalian taxa. Here, using eight poeciliid fish species, we test if species with placental structures invest more resources into offspring brain development than species without placental structures. We predict that matrotrophy may entail higher nutrient provisioning rates to the developing embryo than lecithotrophy, resulting in larger brain sizes in offspring of matrotrophic species, and that a relatively larger part of the total brain growth would occur at younger ages (leading to a shallower ontogenetic brain size allometry). We took non-invasive brain size measurements during the first four weeks of life, and compared these to somatic growth measurements. Contrary to our expectations, we did not find any differences in brain size between the two maternal strategies. Furthermore, we did not find any differences in how relative brain size changed over ontogenetic development, between placental and non-placental species. In contrast to the marsupial/placental transition, the species investigated here only exhibit pre-natal provisioning, which may reduce the potential for maternal investment into brain size. Consequently, our results suggest that coevolution between placental structures and juvenile brain size is not a general pattern.

## INTRODUCTION

Larger brains are considered to have benefits in terms of increased cognitive function (Sol 2009; Kotrschal et al. 2013, 2015a; Heldstab et al. 2016), while being costly to produce and maintain (Barton and Capellini 2011; Isler and Van Schaik 2009; Kotrschal et al. 2013). As a consequence, brain size is considered to evolve in a trade-off between energetic investment and cognitively mediated increases in survival (Sol et al. 2009; González-Lagos et al. 2010; Barton and Capellini 2011). However, increased maternal investment may aid offspring in avoiding energetic trade-offs (Räsänen and Kruuk 2007), by providing increased competitive ability and increased survival rates (Miller 1988; Hutchings 1991; Grindstaff et al. 2003; Bashey 2008; Segers & Taborsky 2011; Leips et al. 2013; Pick et al. 2016; Saporito et al. 2019; but see also Kaplan 1992). Maternal traits that increase energetic investment into offspring may also have a direct influence on the size of energetically costly organs, such as the brain (Martin 1996; Barton and Capellini 2011). Maternal energetic investments may therefore influence brain size, with species that invest more into each individual offspring tending to have larger-brains, compared to species with less investment into each offspring (Pagel and Harvey 1988; Weisbecker and Goswami 2010). For example, gestation period and weaning age are positively correlated with adult brain size in precocial mammals (Isler and Van Schaik 2009; Weisbecker and Goswami 2010), while egg size and care duration are positively correlated with brain size in Tanganyikan cichlids (Tsuboi et al. 2015).

The mode of maternal nutrient transfer has been hypothesized to influence the level of investment into offspring (Weisbecker and Goswami 2010; Mull et al. 2011). For example, pre-natal maternal investment can be divided into two distinct modes: lecithotrophy, where resources for the developing embryo are provided by the egg yolk, or matrotrophy, where resources are transferred to the embryo via either oophagy, surface absorption, a placenta, or placenta-analogous structures (Wourms 1981; Blackburn 1992; Pollux et al. 2014). Across taxa there have been several independent evolutionary transitions between lecithotrophy and matrotrophy, with matrotrophy being widely distributed across both vertebrate and invertebrate taxa (Blackburn 1992; Ostrovsky 2013). In matrotrophic species, resources are continuously supplied to the offspring throughout embryonic development whereas lecithotrophic species are obligated to provide all necessary resources for offspring development prior to fertilization. A direct consequence of these two approaches is that lecithotrophic offspring will lose mass during later embryonic developmental periods, as a cost of metabolism (Thibault and Schultz 1978; Mull et al. 2011; Pollux et al. 2014), compared to matrotrophic offspring which increase mass during embryonic development. Consequently, matrotrophy and placental structures in particular, has likely had a major influence on the evolution of energetic budgets in developing embryos. Indeed, empirical evidence has demonstrated a link between the evolution of the placenta and brain growth, with both the level of placental development and time of gestation being positively correlated with neonate brain size in placental mammals (Elliot and Crespi 2008; Barton and Capellini 2011; but see also Capellini et al. 2011).

Most of our current knowledge about potential links between placental evolution and brain size is derived from mammals, a group with post-natal parental care. In species that exhibit both pre-natal and post-natal care, similar absolute levels of maternal provisioning can be provided by a different relative emphasis placed on either strategy. For example, while hatchlings of altricial bird species are born with smaller brains compared to hatchlings of precocial species, post-natal maternal provisioning increases their relative brain size over ontogenetic development, resulting in larger brains as adults (Iwaniuk and Nelson 2003). Likewise, marsupial mammals are born in a relatively undeveloped state, with brains that are less mature than brains of fetuses of placental mammals at a comparable size (Smith 1997). However, the extended lactation period that occurs in marsupials, enables the development of an adult brain size that is comparable to placental mammals (Weisbecker and Goswami 2010). While post-natal care appears to be highly beneficial for development, many species do not engage in post-natal care. There is limited information about what consequences placental evolution has had on brain size in the large number of taxa that do not exhibit such post-natal parental care. Although, one study on cartilaginous fishes does suggest that matrotrophic species have larger brains than lecithotrophic species (Mull et al. 2011).

Here, we investigate the effects of placental evolution on relative brain size, and ontogenetic brain growth in eight poeciliid fish species (Cyprinodontiformes, Poeciliidae). Importantly, this system lacks post-natal parental care, but does contain several evolutionary transitions between modes of maternal provisioning, i.e between strictly lecithotrophic species with only pre-fertilization provisioning, and highly matrotrophic species with mainly post-fertilization maternal provisioning (Pollux et al. 2014). We use four brain-size measures, to compare between the two maternal strategies during the first four weeks of life. Our prediction was that matrotrophic species will produce offspring with larger brains compared to lecithotrophic species, as matrotrophy may require higher provisioning rates by females. In addition, we hypothesize that species with a smaller initial brain size may show faster brain growth rates, in order to meet potentially similar ecological requirements for the brain size.

## MATERIALS & METHODS

### Study organisms

We used three matrotrophic and five lecithotrophic poeciliid fish species (Table S1, order Cyprinodontiformes; matrotrophic: *Heterandria formosa, Phalloceros caudimaculatus, Poecilia parae*, lecithotrophic: *Gambusia affinis, Girardinus metallicus, Poecilia mexicana, Poecilia reticulata*, and *Xiphophorus nezahualcoyotl)*. According to the latest phylogeny by Furness et al. (2019), each matrotrophic species used in this study represents a separate evolutionary transition to this provisioning mode, with all of the species being situated in distinct clades. For each species, we obtained 12 young adult, captive-bred individuals from the aquarium trade or from laboratory stocks. All experimental procedures were approved by the Ethical Committee in Stockholm, Sweden (license N132/15).

### Data collection

Fish were kept in 13-L tanks and allowed to breed in groups of 4-6 individuals (see supplementary information 1 for details on husbandry). Gravid females with clearly visible embryo-filled abdomens were transferred to separate tanks, and inspected daily for presence of fry. We obtained 4 – 25 fry per species, derived from 1 – 5 females (in total 134 fry, Table S1, supplementary information). The fry were photographed within 24 hours from birth, and then four more times, approximately every seven days, with a digital camera (Canon EOS 1200D; lens: Canon EF 100mm f/2.8 Macro USM; Canon Inc., Tokyo, Japan). The camera was mounted vertically on a copy stand (method modified from Näslund 2014), above a LED light pad (Junlon TRA0609AL; HK Junlon Electronic Technology Co., Ltd, Zhongshan, China), where a small water-containing groove allowed fry to swim in a vertical position. A reference scale was placed beside the fish. Illuminating the fish from below made the brain visible through the semi-transparent skin and cranium. Brain measures derived from this method have previously been found to correlate strongly to measures made from dissected brains (Näslund 2014). Brain and also body size measurements were taken from photographs using ImageJ (Schneider et al. 2012). Specifically, we measured standard body length (from the tip of the nose to the end of the caudal peduncle, i.e. not including the caudal fin) and the brain structures that were clearly visible: width of both optic tecta, length of the right optic tectum, length of the right telencephalon, and width of cerebellum. The telencephalon and cerebellum however were only measured twice, from the first two weeks, as the outline of these structures became difficult to detect over ontogeny.

We used the matrotrophy index from Pollux et al. (2014), defined as the “estimated dry mass of the offspring at birth divided by the dry mass of the egg at fertilization” to assess matrotrophy in our species (see Pollux et al. 2014 for more details). As our species had a rather bimodal distribution in their scores on the matrotrophy index, we used the matrotrophy index as a two-level factor variable, henceforth called “maternal strategy”, which is a measure of whether females do or do not provide offspring with nutrients. A species was categorized as being lecithotrophic when ln-transformed matrotrophy index < 0 (i.e. losing mass over development), and as being matrotrophic when ln-transformed matrotrophy index > 0 (i.e. gaining mass over development).

### Statistical analysis

In order to assess if relative brain size differed between the two maternal strategies, we modelled our log_10_ brain component measures as response variables, with maternal strategy, mean-centered log_10_ transformed body length, and their interaction included as fixed effects. Individual identity and species were included as random effects, allowing for random slopes for species. The four brain measurements (optic tectum width, optic tectum length, telencephalon length, and cerebellum width) where analyzed in separate models.

Further, as the relative growth of the brain depends on the fraction of total somatic growth that is allocated to brain growth, we also assessed whether growth rate differs among the species. To do this, we modeled body length as a response variable and age as a fixed effect, with species and individual identity as random effects. We compared the DIC values of this model to a model that included random slopes for species (representing different growth rates across species). A better fit (lower DIC value) with random slopes suggested among species differences in growth rate.

To explicitly test if growth rate differed between the two maternal strategies, we modeled body length as a response variable, and maternal strategy, mean-centered age, and their interaction as fixed effects. Individual identity and species were fitted as random effects, allowing for random slopes for the species effect.

Finally, it is possible that brain size could trade-off with offspring number. In order to explore if investment into offspring number differed between the maternal strategies, we calculated the species-specific reproductive output per female per day, by dividing the total number of offspring produced during the experiment for each species, by the number of females and by breeding period (in days). The difference in reproductive output of the two maternal strategies was then analyzed using a 2-sample Welch’s *t*-test.

We used Bayesian linear modelling within the MCMCglmm package (Hadfield 2010), in R 3.6.0 to run all models (R Development Core Team 2015). All models were fitted using Gaussian error distributions, an assumption confirmed by residual plots. We chose non-informative priors; a flat, Gaussian prior for the fixed effects, and locally non-informative, inverse Wishart prior for the random effects (following Hadfield 2012). We run 1·10^6^ iterations, a burn-in of 5·10^4^, and a thinning interval of 1000, which resulted in effective posterior sample of 1000 iterations. We diagnosed posterior distributions and model convergence, from three parallel chains, using the Gelman–Rubin convergence criterion; the upper 97.5 quantile of the Gelman-Rubin test statistic was < 1.1 in all cases. All autocorrelations were within the interval −0.1 and 0.1.

The analyses described above did not control for phylogeny, as we had a limited sample of species. However, we note that all matrotrophic species that were included in the study belong to evolutionarily independent transitions. Moreover, while we analyzed matrotrophy as a factorial variable with two levels (placental function, yes/no), all analysis, as described below, were also performed on the continuous matrotrophy index, which gave very similar results (not presented).

## RESULTS

### Growth rate

There were statistical differences in growth rates among the species, as the random species slope model with body length as a response variable and age as a fixed effect represented a better fit to the data than the fixed slope model (DIC_random slope_ = 620, DIC_fixed slope_ = 1265, Fig. 1). There were significant differences in growth rate between the maternal strategies, with matrotrophic species generally increasing less in length than lecithotrophic species over the four week period (β = -0.066; 95% credibility interval (CI): -0.13, -0.015; *P*_MCMC_ = 0.032; Fig. 1, Table S2). In general, individuals of matrotrophic species also tended to be smaller than individuals of lecithotrophic species, although these differences were statistically non-significant (β = -2.27; 95% CI: -4.34, 0.13; *P*_MCMC_ = 0.062; Fig. 1, Table S2).

**Fig 1.**
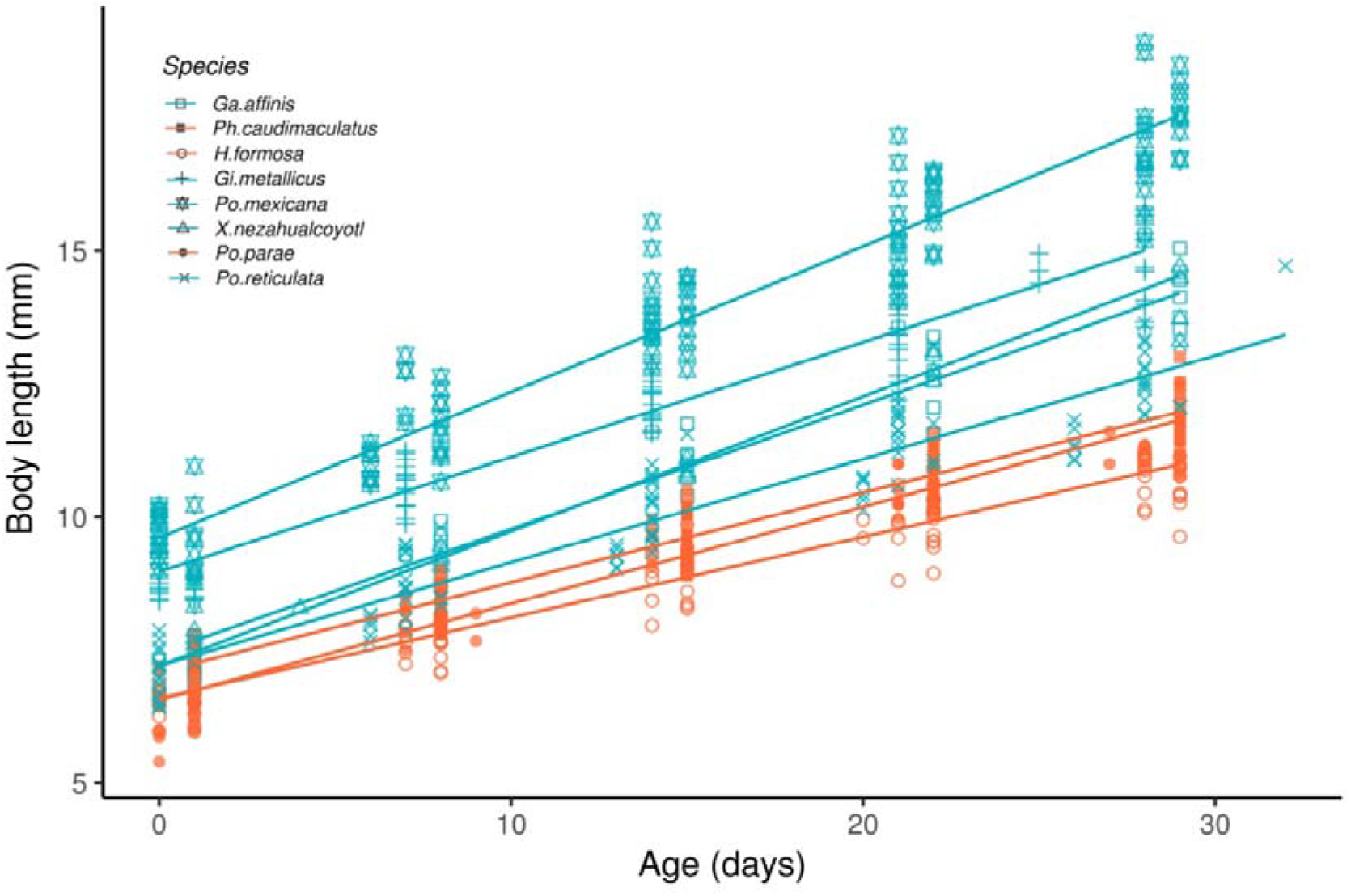
Species-specific slopes of body length (mm) to fish age (days) for lecithotrophic (blue) and matrotrophic (red) species.

### Brain size

We did not find any significant main or interactive effects of maternal strategy (Table 1, Fig 2a-d, Tables S3 – S6), on any of the brain measurements (Table 1, Fig 2a-d, Tables S3 – S6), indicating that there were no clear differences in relative brain size or brain growth across the two maternal strategies, during the first four weeks of life. In three out of four brain measurements, species with a lecitotrophic strategy tended to have larger relative brain sizes, indicating that the relative brain size was, if anything, smaller in matrotrophic species (Table 1; Fig 2).

**Table 1.** Results of brain allometry models.

| **Main effect of Matrotrophy** |  |  |  |  |
| --- | --- | --- | --- | --- |
|  | Mean | Lower 95% CI | Upper 95% CI | P<sub>MCMC</sub> |
| Optic tectum width | -0.036 | -0.077 | 0.01 | 0.1 |
| Optic tectum length | -0.039 | -0.12 | 0.034 | 0.24 |
| Telencephalon | 0.0063 | -0.054 | 0.07 | 0.86 |
| Cerebellum | -0.0084 | -0.049 | 0.026 | 0.59 |
| **Interaction: Matrotrophy X Body Length** |  |  |  |  |
|  | Mean | Lower 95% CI | Upper 95% CI | P<sub>MCMC</sub> |
| Optic tectum width:Body Length | -0.022 | -0.076 | 0.025 | 0.32 |
| Optic tectum length:Body Length | -0.032 | -0.16 | 0.071 | 0.46 |
| Telencephalon:Body Length | -0.051 | -0.44 | 0.34 | 0.78 |
| Cerebellum:Body Length | 0.054 | -0.14 | 0.21 | 0.49 |

Mean=mean value derived from the posterior distribution 95% CI=credibility interval derived from the posterior distribution p<sub>MCMC</sub>=significance value, p<sub>MCMC</sub><0.05 is considered as statistically significant

**Fig 2.**
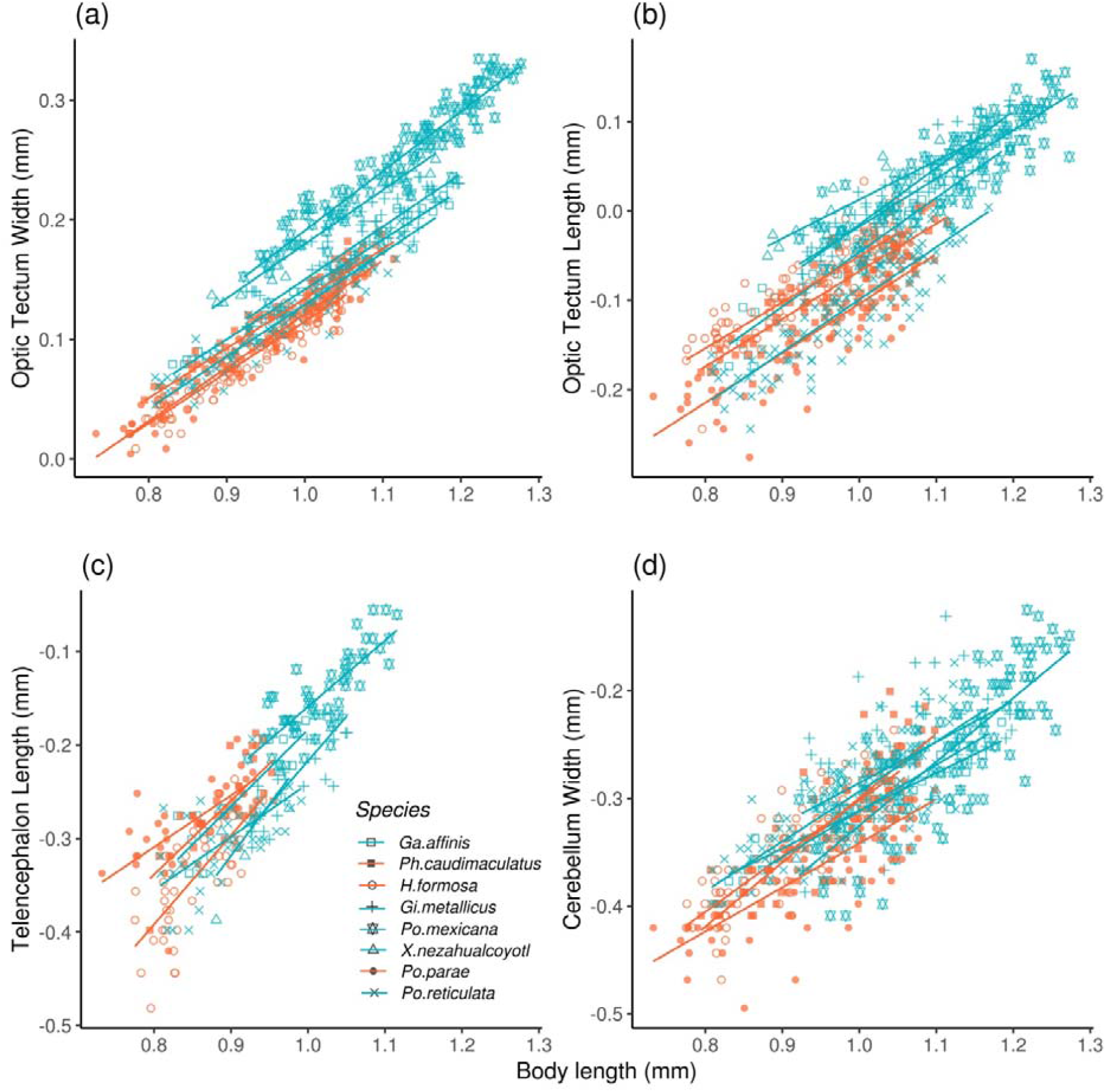
Species-specific allometric slopes of Log_10_-transformed brain measurements to Log_10_-transformed body length (mm): a) optic tectum width, b) optic tectum length, c) telencephalon length, d) cerebellum width. Maternal strategies are represented by blue (lecithotrophic) and red (matrotrophic) colors.

### Reproductive output

There was no significant difference in reproductive output between the two maternal strategies (*t* = -1.16; df = 2.16; *p* = 0.36; 95% CI: -0.51, 0.28; mean_lecithotrophic_ = 0.12; mean_matrotrophic_ = 0.24, Table 1S).

## DISCUSSION

We tested if the evolution of placental structures were associated with increased relative brain size in poeciliid fish. Matrotrophic species provide embryos with nutrients during the whole developmental period, while embryonic development in lecithotrophic species is likely under stronger energetic constraints. We therefore predicted that matrotrophic species would be capable of investing more resources into offspring brain size than lecithotrophic species, but we found no significant differences between the maternal strategies. Instead, our evidence suggested that matrotrophic species tended to have smaller brains, and that the general prediction that matrotrophic species invest more into brain size during embryonic development may not be widespread across taxa.

Previously, investigations into the relationship between the evolution of the placenta/placental structures and brain size have largely been confined to mammals. In mammals, maternal investment via placental nutrition transfer is positively correlated with brain size (Elliot and Crespi 2008; Weisbecker and Goswami 2010; Barton and Capellini 2011). However, differences between groups with and without placental nutrition transfer (i.e. placental and marsupial mammals) decreases with ontogenetic development, presumably because marsupial mammals have a longer lactation period (Weisbecker and Goswami 2010). To date, relatively few studies have explored the link between brain size and placental evolution in taxa that lack post-natal maternal provisioning such as lactation. One study, Mull et al. (2011), did find that placental evolution was positively correlated with relative brain size in small shark species. In contrast, our results do not show the same pattern in poeciliids, potentially because poeciliids differ from sharks in terms of placental function or ecology. In addition, as the study of Mull et al. (2011) examined adults, it is possible that the positive relationship between brain size and matrotrophy in small shark species depends on unmeasured variables that covary with matrotrophy.

In mammals, metabolic rates have also been found to correlate with brain sizes across placental mammals, but not across marsupials (Weisbecker and Goswami 2010). Possibly because the level of placental nutrient transfer is dependent on the energy budget of the female, with the mammalian placenta providing the fetus with direct access to nutrient rich blood, fueling brain development (Weisbecker and Goswami 2010). In poeciliids however, the placenta or placental structure may not be advanced enough to facilitate the level of connection between the mother and embryo, in comparison to many mammals. Additionally, access to fatty acids has also been proposed as a factor promoting rapid brain development *in utero* (Elliot and Crespi 2008). Yet, in matrotrophic poeciliids, nutrient transfer between the mother and developing embryo occurs through epithelial tissues (Scrimshaw 1944), analogous with less derived mammalian placenta types that do not facilitate the transfer of beneficial fatty acids (Scrimshaw 1945, Elliot and Crespi 2008). Mammals and fish also differ substantially in respect to brain development, where in mammals, rapid brain growth occurs when maternal provisioning is high. For example, the brain of placental mammals tends to grow in the fetus, while in marsupials growth occurs during the extended lactation period (Weisbecker and Goswami 2010). Consequently, the mammalian brain develops rapidly in the beginning of life, followed by a much shallower brain allometry after reaching adulthood (Tsuboi et al. 2018). In comparison, the fish brain has continuous growth throughout ontogeny (Tsuboi et al. 2018), possibly meaning that maternal provisioning at the beginning of life is less important for brain development in fishes, compared to mammals.

There are also potentially important ecological differences between matrotrophic and lecitrophic species, which may have contributed to the pattern we observed. For instance, matrotrophy in poeciliids is argued to have evolved under comparatively benign conditions, such as low competition and steady food availability (Trexler and DeAngelis 2003; Pollux and Reznick 2011; Riesch et al. 2012; Tobler and Culumber 2019), which are also associated with increased female fecundity (Thibault and Schultz 1978; Trexler 1997; Trexler and deAngelis 2003, Tobler and Culumber 2019). These relatively benign ecological conditions could also influence selection on brain size and shape among-species brain size differences observed in this poeciliid system. For example, competitive ability and prey capture require advanced cognition, visual, and motor systems (Kaminski et al. 2006; Lankheet et al. 2018; van der Bijl et al. 2018). Therefore, selection for increased brain size may be more relaxed in matrotrophic species, where any benefits of increased cognitive ability are outweighed by high resource availability.

The evolution of placental functions in poeciliids is also likely to be linked to changes in sexual selection. For example, in poeciliids matrotrophy is associated with postzygotic mate choice, and lecithotrophy with prezygotic mate choice (Pollux et al. 2014). In poeciliids, prezygotic mate choice occurs when females inspect male exaggerated secondary sexual traits, such as bright male coloration, courtship behaviors, and ornamental displays (Pollux et al. 2014). Indeed, males from matrotrophic poeciliids species, which are instead more likely influenced by postzygotic mate choice, do tend to have less or lower levels of these secondary sexual traits (Pollux et al. 2014). Mate assessment or choice can be a cognitively demanding process, and therefore any prezygotic sexual selection may also influence selection for increased female brain size (Corral-López et al. 2017). Likewise, the reduced importance of prezygotic mate choice in matrotrophic poeciliids could result in relaxed selection on female cognitive abilities, and consequently brain size. On the other hand, sexual conflict in the form of coercive mating, which is predominantly found in matrotrophic poeciliid species (Pollux et al. 2014), could be expected to lead to selection for increased female brain size. Specifically, as increased brain size and corresponding increased cognitive ability could aid females in avoiding costly matings and aggressive male behavior (Buechel et al. 2016). However, while we found that matrotrophic species instead tended to have smaller brains, overall differences in the pattern of sexual selection across species with and without placentas could offer an explanation why we did not observe any significant differences in brain size.

A trade-off between investment into offspring number and each individual offspring could also affect offspring brain size (Kotrschal et al 2013). In our study, we could only derive a relatively crude measure of female reproductive output. For example, it is possible that some offspring were eaten by females before they were collected, or that our homogeneous laboratory conditions were more favorable to some species compared to others, in the context of reproduction. Therefore, future studies could conduct a more explicit investigation of reproductive output, for example, by counting the number of embryos in females *in vivo*, to investigate if there are difference in maternal investment into somatic tissue between the two maternal strategies. Plastic responses could have also influenced the patterns of maternal investment into brain size that we observed (Mousseau and Fox 1998). Hence, we cannot directly rule out that females, under our laboratory conditions, diverted resources from offspring brain size into other traits, such as offspring somatic growth, or their own reproduction. Although, while the parental generation fish were obtained from several sources, most of the breeding activity was preceded by several weeks of standardized laboratory conditions, which should minimize any influence of plastic parental effects. Finally, we also acknowledge that examining brain size on a scale that is relative to body size, opens up the risk of divergences in body size across taxa leading to inferential errors (see Rogell et al. 2020). We consider this unlikely however, as while matrotrophic species were smaller than lecithotrophic species, relative brain sizes were similar, suggesting that inferential errors are not problematic in our study.

In conclusion, we found no relationship between offspring brain size and maternal investment strategy (matrotrophic or lecithotrophic) in poeciliid fish. In contrast, previous studies on mammals and cartilaginous fishes have demonstrated that brain size can be associated with the level of placental development. Therefore, our results suggest that the link between placental evolution and brain size is not necessarily a general pattern across taxa as previous predicted. Instead, we suggest that the placenta or placental structures in poeciliids are not as sufficiently developed as in other taxa, that ecological differences among-species may have obscured expected patterns, or that the specific ontogenetic development of the fish brain does not produce a large brain size during early life stages.

The experimental procedures were approved by the Ethical Committee in Stockholm, Sweden (license N132/15).

The authors declare no conflicts of interests.

Data will be deposited to the Dryad archive together with article publication.

## Acknowledgements

We would like to thank Niclas Kolm and John Fitzpatrick for providing some of the fish used in this experiment. We thank Raïssa de Boer for help with experimental procedures, and Bertil Borg for constructive comments on the manuscript. This research was supported by The Swedish Research Council grant to B.R. (grant number 2013-05064). The authors declare no conflicts of interests. The experimental procedures were approved by the Ethical Committee in Stockholm, Sweden (license N132/15).

