## Supplementary information for "The evolution of a placenta is not linked to increased brain size in poeciliid fishes"

**Supplementary information 1:** Experimental procedures

Each group tank contained gravel bottom, small sponge filter, an empty flower pot and a wool yarn mop as a shelter. The fish were fed to satiation three times a day (one time a day during weekends) with a mixture of freshly hatched artemia naupli and defrosted chironomid larvae. Water was changed once a week. 30% of aquarium water was replaced with tap water with addition of 1.11g marine aquarium salt per 1 liter of water (pH=7.6, Carbonate hardness=10°dKH, General hardness=20°dH). The fry were kept individually in 1 liter tanks, fed with artemia naupli to satiation three times a day (one time a day during weekends), and 80% of water was changed three times per week. The tanks were placed in a temperature-controlled rum, resulting in water temperatures of 25°C.

Table S1. Species list with details on maternal strategy and number of families and individuals used in the study.

| **Species** | **ln(MI*)** | **maternal strategy** | **nr. families** | **nr. individuals** | **reproductive output^†^** |
| --- | --- | --- | --- | --- | --- |
| *Gambusia affinis* | -0,48 | lecithotrophic | 3 | 8 | 0,116 |
| *Girardinus metallicus* | -0,33 | lecithotrophic | 3 | 17 | 0,055 |
| *Poecilia mexicana* | -0,47 | lecithotrophic | 3 | 25 | 0,118 |
| *Poecilia reticulata* | -0,41 | lecithotrophic | 6 | 23 | 0,143 |
| *Xiphophorus nezahualcoyotl* | -0,67 | lecithotrophic | 1 | 4 | 0,174 |
| *Heterandria formosa* | 3,55 | matrotrophic | 5 | 24 | 0,163 |
| *Phalloceros caudimaculatus* | 0,76 | matrotrophic | 2 | 14 | 0,429 |
| *Poecilia parae* | 1,91 | matrotrophic | 5 | 19 | 0,117 |

*Matrotrophy index

^†^Total reproductive output: number of offspring per female, per day

**Supplementary information 2:** Model outputs

Table S2. Output of the model investigating growth differences between the maternal strategies.

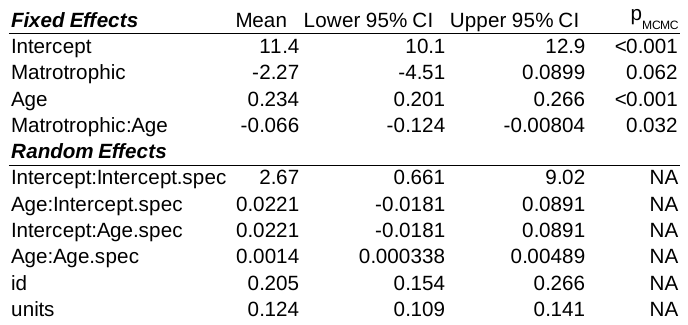

Table S3. Output of the model with **optic tectum width** as a response variable and maternal strategy, body length, and their interaction as fixed effects.

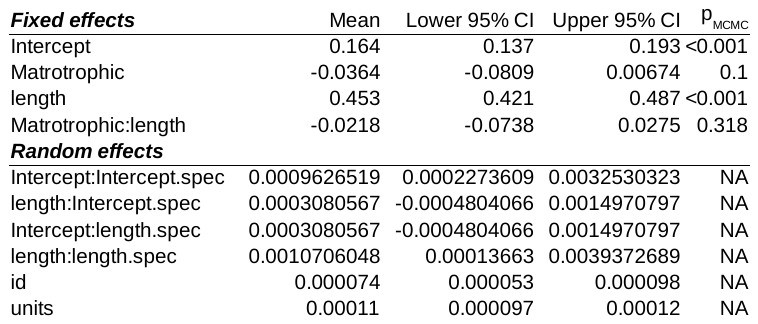

Table S4. Output of the model with **optic tectum length** as a response variable and maternal strategy, body length, and their interaction as fixed effects.

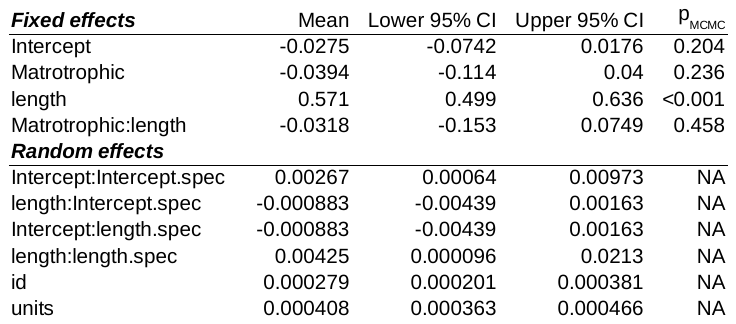

Table S5. Output of the model with **telencephalon** length as a response variable and maternal strategy, body length, and their interaction as fixed effects.

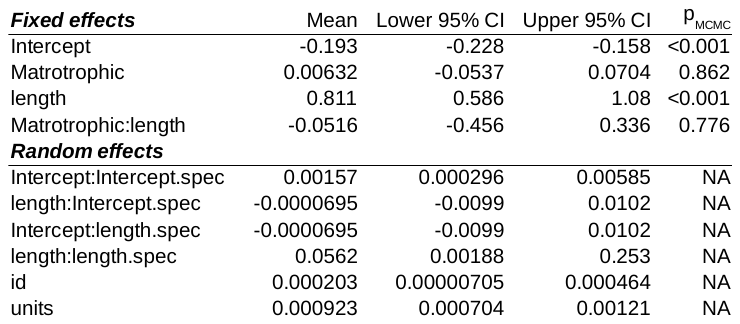

Table S6. Output of the model with **cerebellum** width as a response variable and maternal strategy, body length, and their interaction as fixed effects.

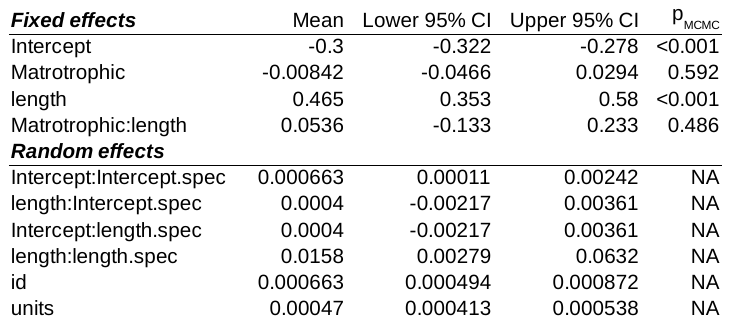
